# Ketone body mediated histone β-hydroxybutyrylation is reno-protective

**DOI:** 10.1101/2024.12.18.628574

**Authors:** Juthika Mandal, Sachin Aryal, Ishan Manandhar, Saroj Chakraborty, Xue Mei, Beng San Yeoh, Blair Mell, Andrew Kleinhenz, Ramakumar Tummala, Tao Yang, Piu Saha, William T Gunning, Matam Vijay-Kumar, Venkatesha Basrur, Ivana de la Serna, Bina Joe

## Abstract

Starvation, intermittent fasting and exercise, all of which are recommended lifestyle modifiers share a common metabolic signature, ketogenesis to generate the ketone bodies, predominantly β-hydroxybutyrate. β-hydroxybutyrate exerts beneficial effects across various contexts, preventing or mitigating disease. We hypothesized that these dynamic health benefits of β-hydroxybutyrate might stem from its ability to regulate genome architecture through chromatin remodeling via histone β-hydroxybutyrylation, thereby influencing the transcriptome. Focusing on the kidney, which is an end organ protected by β-hydroxybutyrate, we examined histone β-hydroxybutyrylation-mediated chromatin remodeling. Notably, regions of the genome associated with lipid catabolism were predominantly in an open chromatin configuration, leading to active transcription and translation. Significant β-hydroxybutyrylation was observed in the kidneys and the most highly upregulated gene actively transcribed and translated was 3-hydroxy-3-methyglutaryl CoA Synthase 2 (*Hmgcs2*), a gene responsible for the biosynthesis of β-hydroxybutyrate in mitochondria. In contrast, regions with more compact chromatin structures were enriched with genes related to immune function such as protein tyrosine phosphatase receptor type C (*Ptprc*) and lymphocyte cytosolic protein 1 (*Lcp1*), which exhibited reduced transcription and translation. These results reveal that renal epigenetic histone β-hydroxybutyrylation is a novel mechanism by which transcriptional regulation of both energy metabolism and immune function occur concomitantly to protect kidneys and lower hypertension.

Graphical Abstract

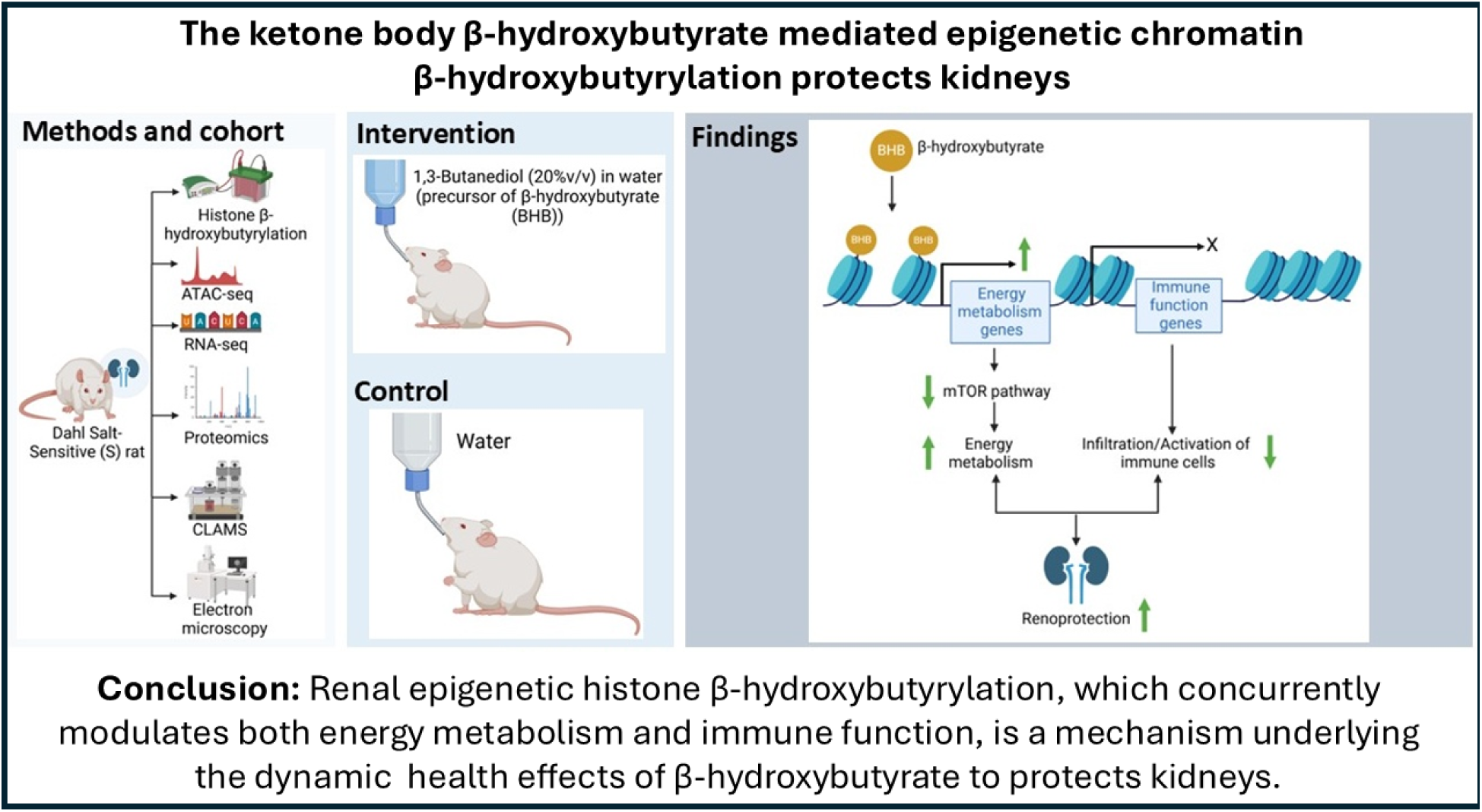

## Introduction

β-hydroxybutyrate is a ketone body produced in the liver during the breakdown of fatty acids during low carbohydrate intake, fasting, or exercise^1^. It is a crucial alternate energy substrate which makes β-hydroxybutyrate essential during states of ketosis, where the body shifts from using carbohydrates to using fats for energy^1,2^. β-hydroxybutyrate is not only a fuel; it also functions as a signaling molecule that can influence gene expression^2^. It modulates pathways related to energy metabolism, inflammation, and oxidative stress^3–5^. β-hydroxybutyrate is shown to inhibit histone deacetylases, which affect gene transcription, potentially promoting longevity and reducing inflammation^4,6,7^. Particularly related to inflammation, β-hydroxybutyrate is reported to downregulate the expression of the NLRP3 inflammasome^7–9^. Due to all of these known beneficial effects, there is a growing interest in leveraging β-hydroxybutyrate to help manage chronic metabolic diseases, such as type 2 diabetes, obesity, and cardiovascular diseases^3,9–13^.

Metabolism plays a key role in hypertension. Previously, we and others reported metabolic perturbations in salt-sensitive hypertension (PMID: 32336227, 19546378, 30595123). Among them, β-hydroxybutyrate is associated with lower blood pressure and kidney damage. Next, we examined the effect of reconstituting β-hydroxybutyrate in the Dahl rat and discovered that it has a profound antihypertensive effect, which was associated with a betterment of renal function^9^. However, the molecular mechanism by which β-hydroxybutyrate protects kidneys remains unknown.

Here we focused on the kidney and specifically examined whether the mechanism underlying the beneficial effect of β-hydroxybutyrate was due to β-hydroxybutyrylation, which is a newer epigenetic histone modification caused by β-hydroxybutyrate. Data in support of histone β-hydroxybutyrylation mediated epitranscriptional regulation of metabolism and immunity as a novel reno-protective mechanism are presented.

## Methods

### Monitoring Energy Metabolism and activity

Metabolic parameters were measured in 12 to 13-week-old male rats after 8 weeks of 1,3-butanediol supplementation. Rats were housed individually for 24 hrs in the Comprehensive Laboratory Animal Monitoring System (CLAMS, Columbus Instruments). Oxygen consumption (VO_2_) and carbon dioxide production (VCO_2_) were sampled sequentially for 5 seconds in a 10-minute interval and motor activity was recorded every second in X and Z dimensions. Respiratory exchange ratio (RER) was calculated as VCO_2_/VO_2_.

### Food Intake

Food given to rats housed in CLAMS was measured before and after 24hrs. The difference was recorded as food intake.

### Measurement of Serum BHB

Serum β-hydroxybutyrate level was measured using a colorimetric assay kit from Cayman Chemicals.

### Histone Extraction

Snap-frozen kidney samples were homogenized for histone extraction using the EpiQuik histone extraction kit and quantitated using the Bicinchoninic Acid method^15^. Proteins were aliquoted and stored at −80°C until further use.

### Western Blotting

Histones were resolved by gel electrophoresis on a 4-20% gradient SDS PAGE gel and probed using primary antibodies from PTM Biolabs (www.ptmbiolabs.com).

### Molecular Analyses

Detailed methods for Molecular Analyses are under Supplementary Methods. Briefly, kidneys were examined for chromatin accessibility via ATAC-seq and Chromatin immunoprecipitation (ChIP) assays, gene expression via RNA-seq and real-time PCR, and protein abundance by mass spectrometry-based quantitative proteomic analysis. Primer sequences are in Tables S1 and S2.

### Pathway Analyses

KEGG based pathway enrichment analyses were conducted using ShinyGOv0.77 (FDR cut off <0.05)^16^. Reactome pathway knowledgebase was used to generate pathways^17^.

### Transmission Electron Microscopy and Analysis of Mitochondrial Morphology

Transmission electron microscopy was used to examine the impact of β-hydroxybutyrylation on mitochondrial morphology in the proximal tubule epithelial cells as detailed under Supplementary Methods.

### Quantification of peripheral T cells proliferation

Peripheral lymphocytes were isolated from blood using the Histopaque method^18^. Briefly, lymphocytes (1×10^6^ cells/well) were stained with carboxyfluorescein diacetate succinimidyl ester (CFSE, Sigma, 5 µM), washed and cultured in 1% heat inactivated fetal bovine serum in DMEM media for 5 days followed by staining with anti-rat CD3-APC antibody for loading onto the flow cytometer (BD AccuriTM C6 Plus).

### Histological Analyses

Kidneys were fixed in neutral buffered formalin for 24 hours and transferred to 70% ethanol for paraffin-embedding and sectioning. Sections were stained with Hematoxylin & eosin. Rabbit anti-PMP70 antibody (ThermoFisher Scientific, Cat# PA1-650, 1:100 dilution) was used and detected by using Vectastain Elite ABC kit and SigmaFast 3,3-Diamino-benzidine tablets. The images were captured by VS120 Virtual Slide Microscope (Olympus) and analyzed using Olyvia Ver.2.9.

### Quantification and Statistical Analyses

GraphPad Prism version 9.3.1 was used for data analysis. Statistical analyses were performed using the student’s t-test or one-way ANOVA with Bonferroni post hoc test as necessary. Data with a p-value < 0.05 was considered significant. All data: Mean ± SEM.

## Results

### 1,3-Butanediol is a precursor of β-hydroxybutyrate in both sexes

Oral administration of 1,3-butanediol to male rats increased their circulating β-hydroxybutyrate^9^. Here we examined whether the ability of 1,3-butanediol to reconstitute serum β-hydroxybutyrate was sex-independent. As seen in Figure 1A, B, both male and female rats responded to 1,3-butanediol treatment by elevation of circulating β-hydroxybutyrate indicating that the conversion of 1,3-butanediol to β-hydroxybutyrate occurred independent of sex.

**Figure 1:**
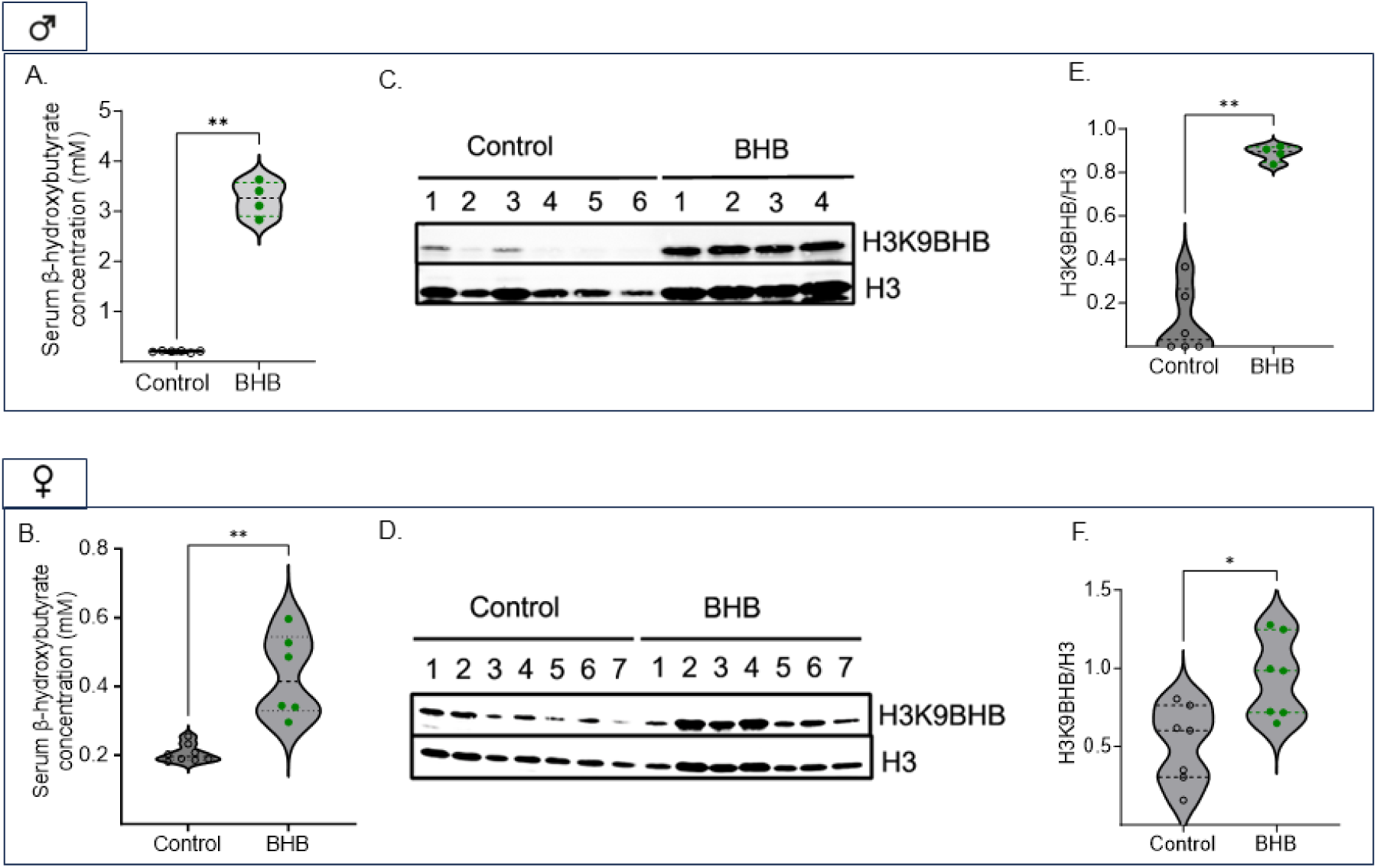
Histone β-hydroxybutyrylation is elevated with 1,3-butanediol treatment. (A-B) Serum levels of β-hydroxybutyrate in males and females. (C-D) Immunoblotting for β-hydroxybutyrylation. (E-F) Quantification of the blots. BHB: β-hydroxybutyrate, H3K9BHB: Histone 3 lysine 9 β-hydroxybutyrylation, H3: Histone 3. All data are mean ± SEM, *p<0.05, **p<0.01.

### Enhanced Histone β-hydroxybutyrylation in 1,3-butanediol treated rats

Next, we investigated if β-hydroxybutyrate resulted in histone 3 lysine-9 β-hydroxybutyrylation, which is the modification reported to be affected during starvation ^2^. As seen in Figures 2C and 2D, both males and females treated with 1,3-butanediol, exhibited a significant increase in renal histone 3 lysine-9 β-hydroxybutyrylation. These data are presented with equal protein loading of isolated histones. Despite this, for reasons unknown, we noticed that the levels of H3 were also elevated in samples from the rats treated with 1,3-butanediol. However, to maintain rigor, β-hydroxybutyrylation was quantified using H3 as the normalizing factor. As seen in Figures 2E and 2F, β-hydroxybutyrylation was more prominent in males than in females. For all subsequent studies, male rats were used. Henceforth, this group of male rats with increased renal histone 3 lysine 9 β-hydroxybutyrylation will be referred to as the BHB group.

**Figure 2:**
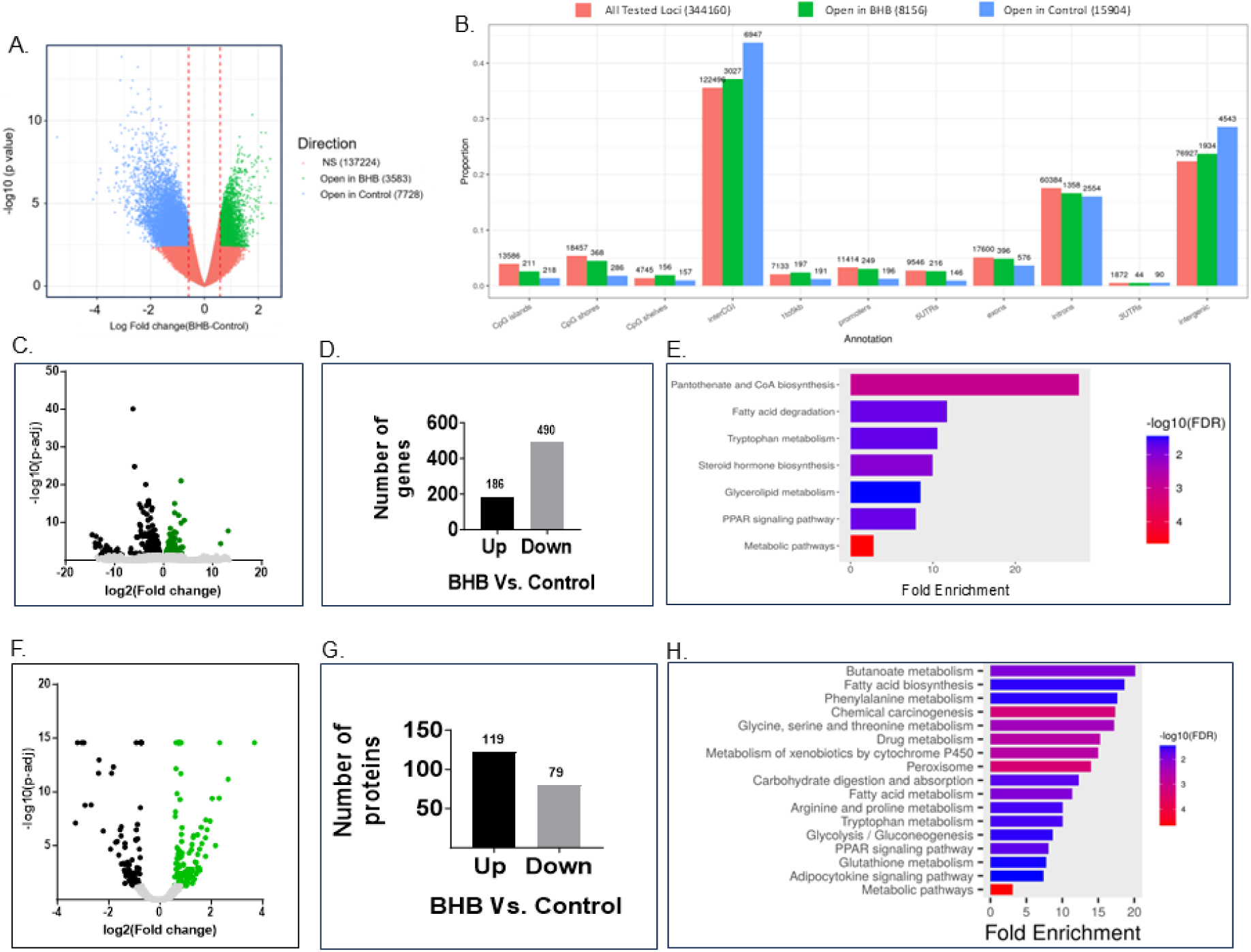
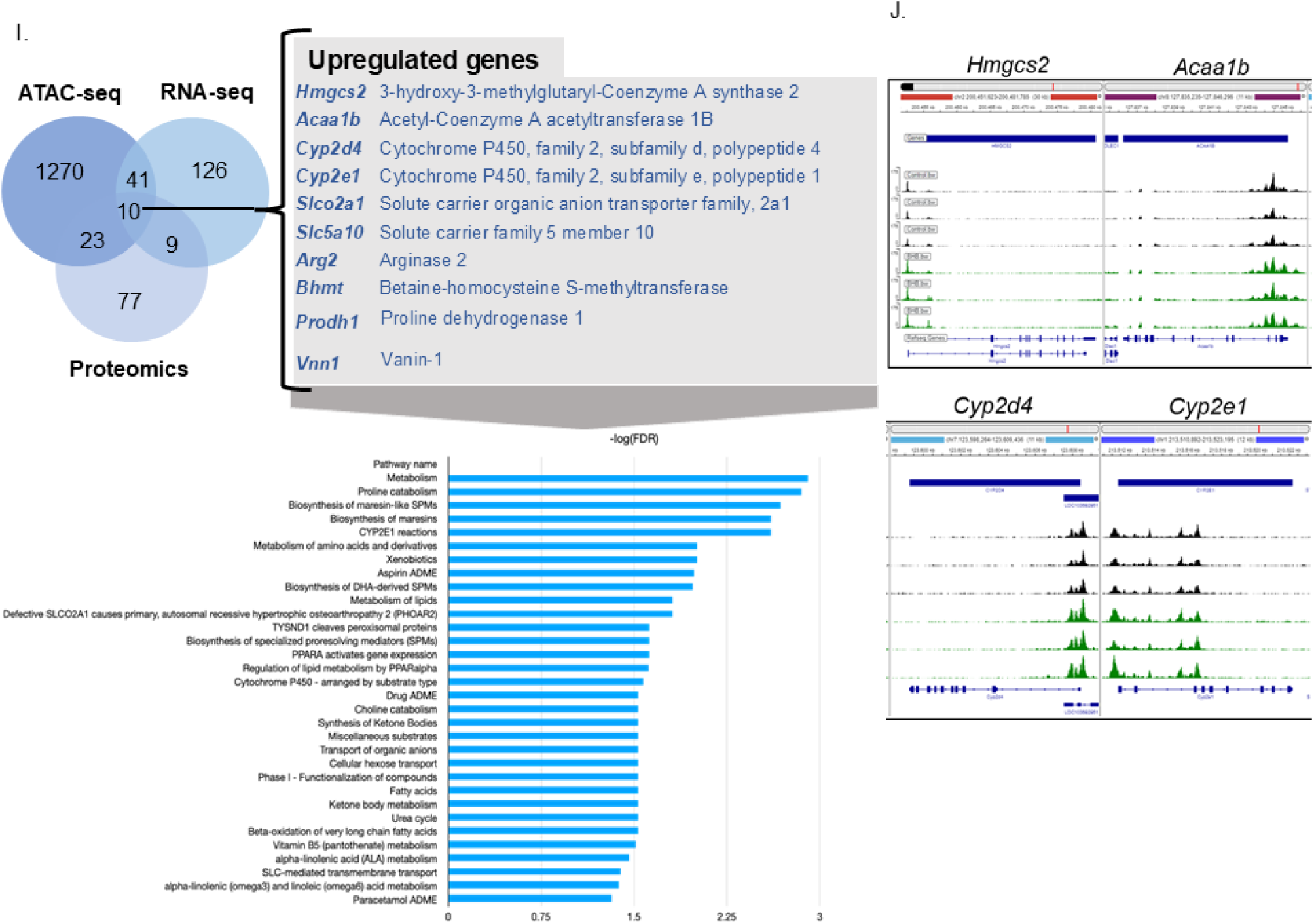
Chromatin accessibility, gene expression, and protein levels are altered with 1,3-butanediol treatment. (A) Volcano plot for BHB vs. control groups. Each data point is a tested locus. The vertical lines correspond to the logFC cutoff. (B) Annotation summary of loci tested in BHB vs control groups. X-axis: genomic annotations; Y-axis proportion of loci in each category within each type of annotation. All Tested Loci (red bars), serve as a basis by which to compare the significant loci (blue and green bars). (C) RNAseq data: Volcano plot representing the significantly differential transcripts. The X-axis: log2 normalized fold change in BHB group compared to control; The Y-axis: – log10 of adjusted p value. (D) Differentially expressed transcripts in numbers. The black bar indicates upregulated transcripts, and the gray bar indicates downregulated transcripts in the BHB group compared to the control. (E) Pathways associated with the upregulated genes of BHB group. (F) Proteomics data: Volcano plot depicting significantly altered protein levels. X-axis: log2 normalized fold change in the BHB group compared to control. Y-axis: –log10 of adjusted p value. (G) Differentially expressed proteins in numbers. The black bar indicates upregulated proteins, and the grey bar indicates downregulated proteins in BHB groups compared to the control. (H) Pathways associated with upregulated proteins in the BHB group. (I) Venn diagram showing common upregulated genes and corresponding proteins leading to increased metabolism. Different shades of blue circles depict ATACseq, RNAseq, and Proteomics data and numbers within the circles are numbers of genes/proteins significantly upregulated. The gray box provides the name for common upregulated genes and proteins; Underneath are the associated upregulated pathways. (J) Chromatin accessibility for genes of interest. The y-axis represents chromatin cut sites and thus open chromatin, and x-axis represents the chromatin location of genes of interest. Black peaks represent the control group, and green peaks represent BHB group.

### Histone β-hydroxybutyrylation promoted large-scale chromatin remodeling

To assess the extent of differential chromatin accessibility attributed to histone β-hydroxybutyrylation, we performed ATAC-sequencing. In the BHB group, 3583 genomic regions were detected in the open configuration implying enhanced accessibility for transcriptional regulation (Figure 2A). Similarly, 7728 loci were ‘open’ in the control group indicating that in the BHB group these regions were ‘closed’ or compacted regions of the genome with decreased accessibility for transcription (Figure 2A). Figure 2B shows the proportions of loci at various genomic locations. Most of the remodeling occurred in the interCGI regions (regions of the genome that lie between CpG islands) followed by intergenic regions (parts of the genome that lie between genes) and introns. Interestingly, 249 and 196 promoter regions were open in the BHB and control groups respectively (Figure 2B).

### Histone β-hydroxybutyrylation-mediated remodeling of the renal transcriptome and proteome

To uncover the extent to which histone β-hydroxybutyrylation regulated the transcriptome, we performed renal RNA-sequencing. In the BHB group compared to the control, a total of 186 genes were upregulated, and 490 genes were downregulated (Figures 3C and 3D). Pathway analysis revealed that the top 2 upregulated pathways were pantothenate and CoA biosynthesis and fatty acid degradation (Figure 2E).

**Figure 3:**
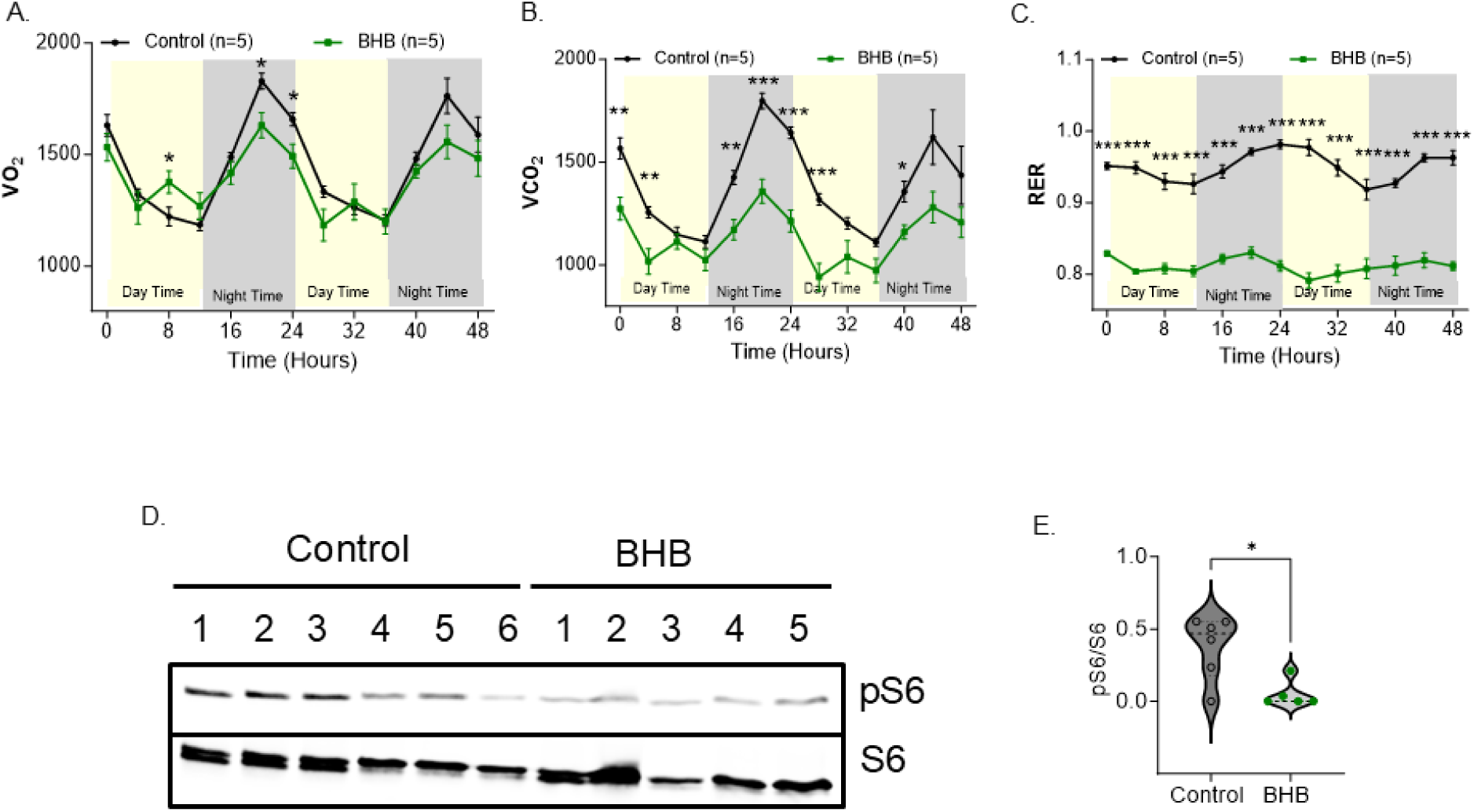
1,3-butanediol treatment reduces respiratory exchange ratio (RER) and inhibits mammalian target of rapamycin complex 1 (mTORC1). (A-C) Metabolic CLAMS data showed reduced VO_2_, VCO_2_, and RER level in the BHB group compared to controls. Black line: control group and green line: BHB group. (D-E) Reduced phospho-S6 ribosomal protein levels were found in male rats treated with BHB compared to control. See also Supplementary Figure S4 for data from female rats. BHB: β-hydroxybutyrate. All data are mean±SEM; *p ≤ 0.05, **p ≤ 0.01, and ***p ≤ 0.001.

Next, to examine the impact of the chromatin-remodeling responsive transcriptome on the proteome, we conducted a mass spectrometry-based quantitative proteomic analysis. A volcano plot of the differential proteome is shown in Figure 2F. A total of 119 and 79 proteins were significantly upregulated and downregulated, respectively, in the BHB group compared to control (Figure 2G). Pathway analysis of the proteomics data indicated that butanoate metabolism, fatty acid biosynthesis, and fatty acid metabolism (Figure 2H) were the upregulated pathways whereas pathways downregulated in the BHB group were all related to immune function (Supplementary Figure 1A and Figure 1B).

### Chromatin remodeling by histone β-hydroxybutyrylation upregulated lipid catabolism

Next, we explored if the observed alterations in the transcriptome and proteome of the BHB group was specifically due to chromatin remodeling via histone β-hydroxybutyrylation. We prioritized common differentially regulated outputs between chromatin states identified by ATAC-Seq, RNA-seq, and proteomics datasets. Such a combinatorial analysis revealed that there were 10 upregulated proteins aligned with enhanced transcription caused by histone β-hydroxybutyrylation (Figure 2I). Intriguingly, four of the top genes share a common function, lipid metabolism. These were 3-Hydroxy-3-methylglutaryl-Coenzyme A synthase 2 (*Hmgcs2*), Acetyl-Coenzyme A acetyltransferase 1B (*Acca1b*), Cytochrome P450, family 2, subfamily d, polypeptide 4 (*Cyp2d4*), Cytochrome P450, family 2, subfamily e, polypeptide 1 (*Cyp2e1*) (Figure 2I). Chromatin accessibility of the promoter regions of *Hmgcs2, Acaa1b, Cyp2d4,* and *Cyp2e1* were higher in the BHB group compared to control (Figure 2J). These data indicated that histone β-hydroxybutyrylation caused open chromatin within the promoter regions of *Hmgcs2, Acaa1b, Cyp2d4,* and *Cyp2e1.* Reactome pathway analysis using the 10 genes identified metabolism as the top upregulated pathway along with metabolism of lipids, synthesis and metabolism of ketone bodies, and β-oxidation of very long chain fatty acids (Figure 2I).

Collectively, these results led us to further prioritize β-hydroxybutyrylation as a key mechanism promoting lipid catabolism as a primary source for energy.

### Metabolic reprogramming in response to elevated histone β-hydroxybutyrylation

To determine if rats treated with 1,3-butanediol are preferring lipids for energy, we housed both control and BHB group of rats in the Comprehensive Lab Animal Monitor System (CLAMS) and monitored their metabolic parameters. Overall, both oxygen consumption (VO_2_) and carbon dioxide production (VCO_2_) were lower in the BHB group compared to control (Figure 3A and 4B). Importantly, respiratory exchange ratio (RER) was dramatically decreased in the BHB group compared to the control group (Figure 3C). A lower RER points to lipids, but not carbohydrates as the primary source of energy. These data provided further evidence for histone β-hydroxybutyrylation enhanced transcription of key metabolic genes in the lipid mobilizing pathways. Those key genes, *Hmgcs2, Acaa1b, Cyp2d4 and Cyp2e*1 likely contributed to a preferential switch in bioenergetic fuel source to lipids (Figure 3C).

**Figure 4:**
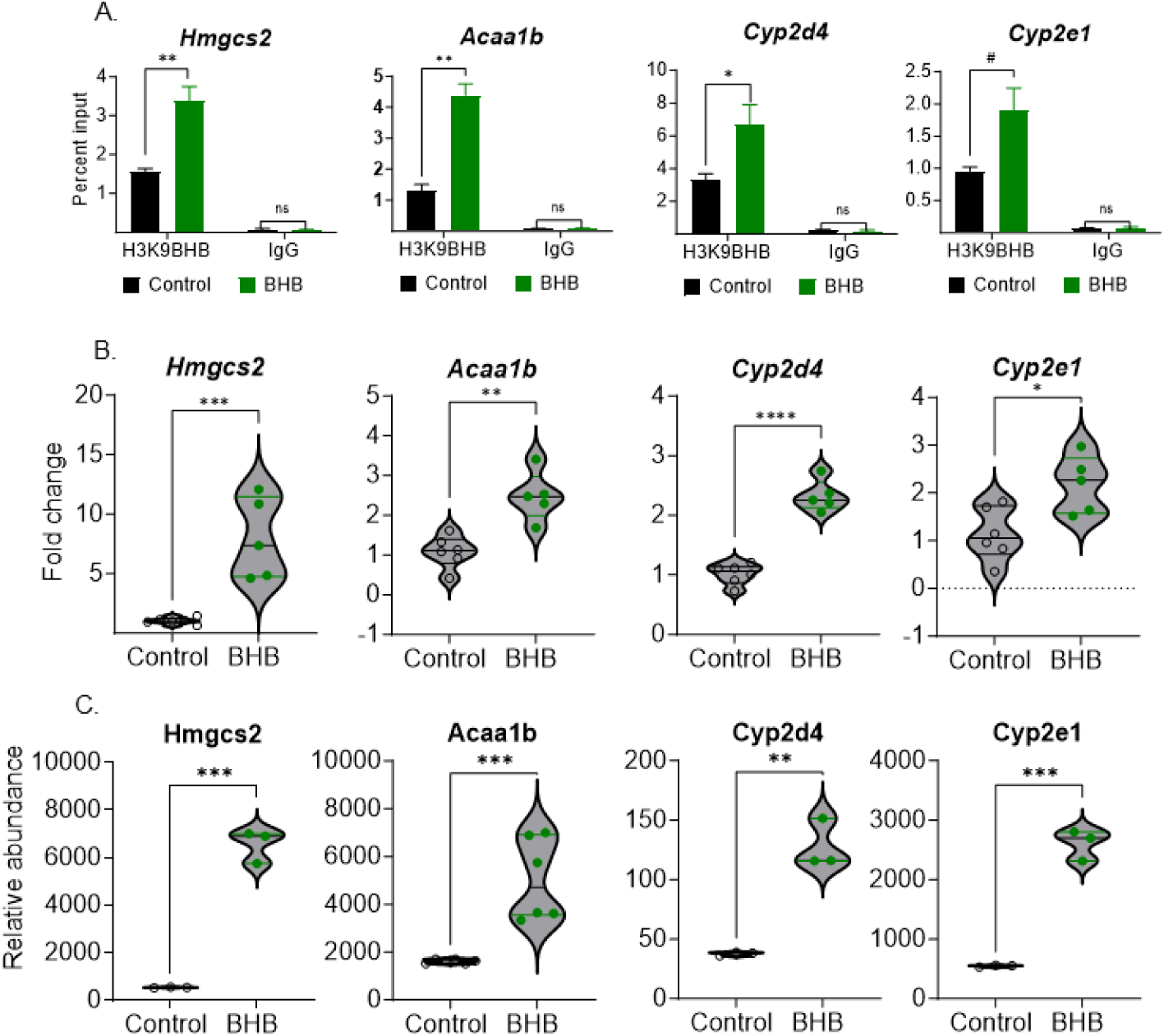
Validation of common upregulated genes through CHIP-qPCR, Real time PCR, and proteomics. (A) CHIP-qPCR data demonstrating enrichment of *Hmgcs2, Acaa1b, Cyp2d4,* and *Cyp2e1* in the BHB group compared to control (n=3 replicates/group). Black bar: Control, Green bar: BHB. (B) Real time PCR showed higher expression of *Hmgcs2*, *Acaa1b*, *Cyp2d4*, and *Cyp2e1* in BHB group compared to control (housekeeping gene is *L36a*). (C) Normalized relative abundance of Hmgcs2, Acaa1b, Cyp2d4, Cyp2e1 proteins. See also Supplementary Figure S2 for female data. Black open circle-control, green closed circle-BHB. All data are mean±SEM; *p< 0.05, **p< 0.01, ***p< 0.001, ****p <0.0001 and ^#^p=0.0567.

### *Hmgcs2*, *Acaa1b*, *Cyp2d4*, and *Cyp2e1* are bonafide targets linking epigenetic renal Histone-3 β-hydroxybutyrylation to energy metabolism

In addition to β-hydroxybutyrylation, β-hydroxybutyrate is also known to epigenetically modify histones by acetylation^7,19^. Therefore, it was important to determine whether the 4 upregulated energy metabolism genes, *Hmgcs2*, *Acaa1b*, *Cyp2d4*, and *Cyp2e1*, which were prioritized in our study were specific targets of β-hydroxybutyrylation. We therefore performed Chromatin immunoprecipitation (ChIP) assays using an antibody specific to β-hydroxybutyryl-histone H3 (Lys 9) β-hydroxybutyrate. ChIP assay using this antibody revealed that promoter regions of *Hmgcs2*, *Acaa1b*, *Cyp2d4*, and *Cyp2e1* were significantly enriched in the BHB group (Figure 4A), thus confirming that these genes were bonafide targets of β-hydroxybutyrylation.

Next, transcripts of these genes were quantitated by real time qPCR analysis. The abundance of Hmgcs2 was prominently upregulated by ∼15 fold higher in the BHB group compared to control (Figure 4B, Supplementary Figure 1A). Similarly, the abundances of *Acaa1b*, *Cyp2d4* and *Cyp2e1* were also upregulated, *albeit* to lower extents of 3-4-fold in the BHB group compared to controls (Figure 4B, Supplementary Figure 1B-2D). Aligned with these data, quantitative mass-spectrometry using high-quality MS3 spectra indicated that all 4 protein products of *Hmgcs2*, *Acaa1b*, *Cyp2d4* and *Cyp2e1* were significantly upregulated in the BHB group compared to control (Figure 4C). Taken together, these data demonstrate that β-hydroxybutyrate, via conferring epigenetic histone β-hydroxybutyrylation, upregulated enzymes which mobilize lipids for energy metabolism.

### Upregulation of Hmgcs2-mediated fatty acid oxidation increased renal proximal tubule epithelial cell mitochondrial circulatory index

One of the prominent targets upregulated due to histone β-hydroxybutyrylation was the mitochondrial protein Hmgcs2. Hmgcs2 is a cytosolic and rate-limiting enzyme regulating mitochondrial fatty acid oxidation to promote ketogenesis. Ketogenesis is largely attributed to the liver, but recent work demonstrates that Hmgcs2 in the kidneys also participates in this pathway^20,21^. Because ketogenesis in the liver increases mitochondrial circularity, we asked if such alterations in the ultrastructure of mitochondria also occur in the kidney. Transmission electron microscopy revealed more circular mitochondria in the proximal tubule epithelial cells from the BHB group compared to the elongated mitochondria in the controls (Figure 5A). This observation was confirmed by morphometric analyses of the measurement of average mitochondrial area, area/perimeter, and circularity index of these mitochondria. As seen in Figures 6B, 6C and 6D, each of these morphometric measures were elevated in the BHB supplemented group compared to the controls. These data support our conclusion that histone β-hydroxybutyrylation mediated upregulation of Hmgcs2 to promote ketosis contributed to enhanced renal mitochondrial circularity index.

**Figure 5:**
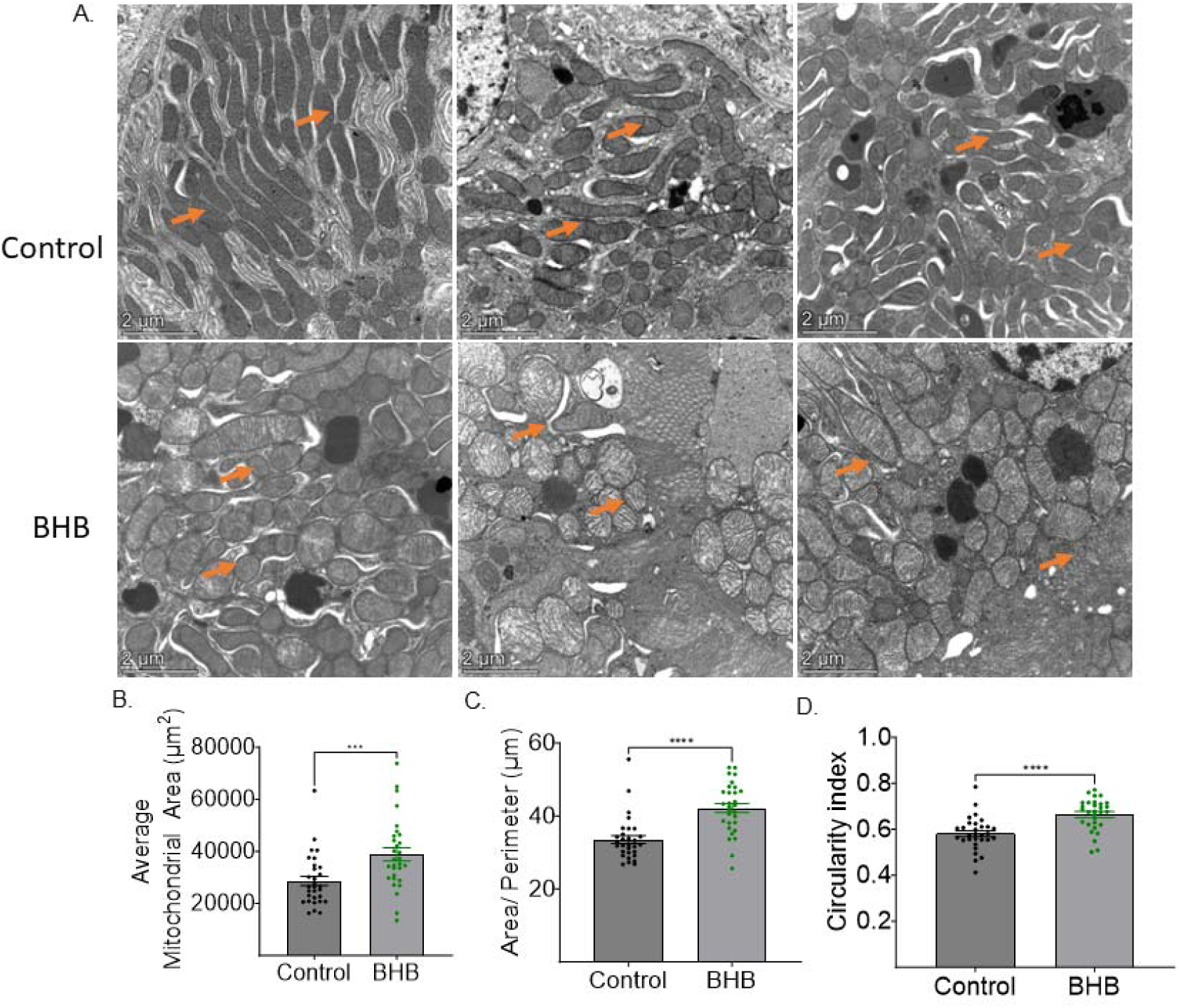
Kidney proximal tubule epithelial cell mitochondrial circulatory index was found to be higher in BHB group compared to control. (A) Representative transmission electron microscopy images of renal mitochondria. Orange arrows point to mitochondria. (B-D) Bar graph shows quantification (B) Average mitochondrial area (C) Area/Perimeter, and (D) Circularity index. BHB: β-hydroxybutyrate. n=30 images per group. All data are mean±SEM, ***p< 0.001, and ****p<0.0001.

**Figure 6:**
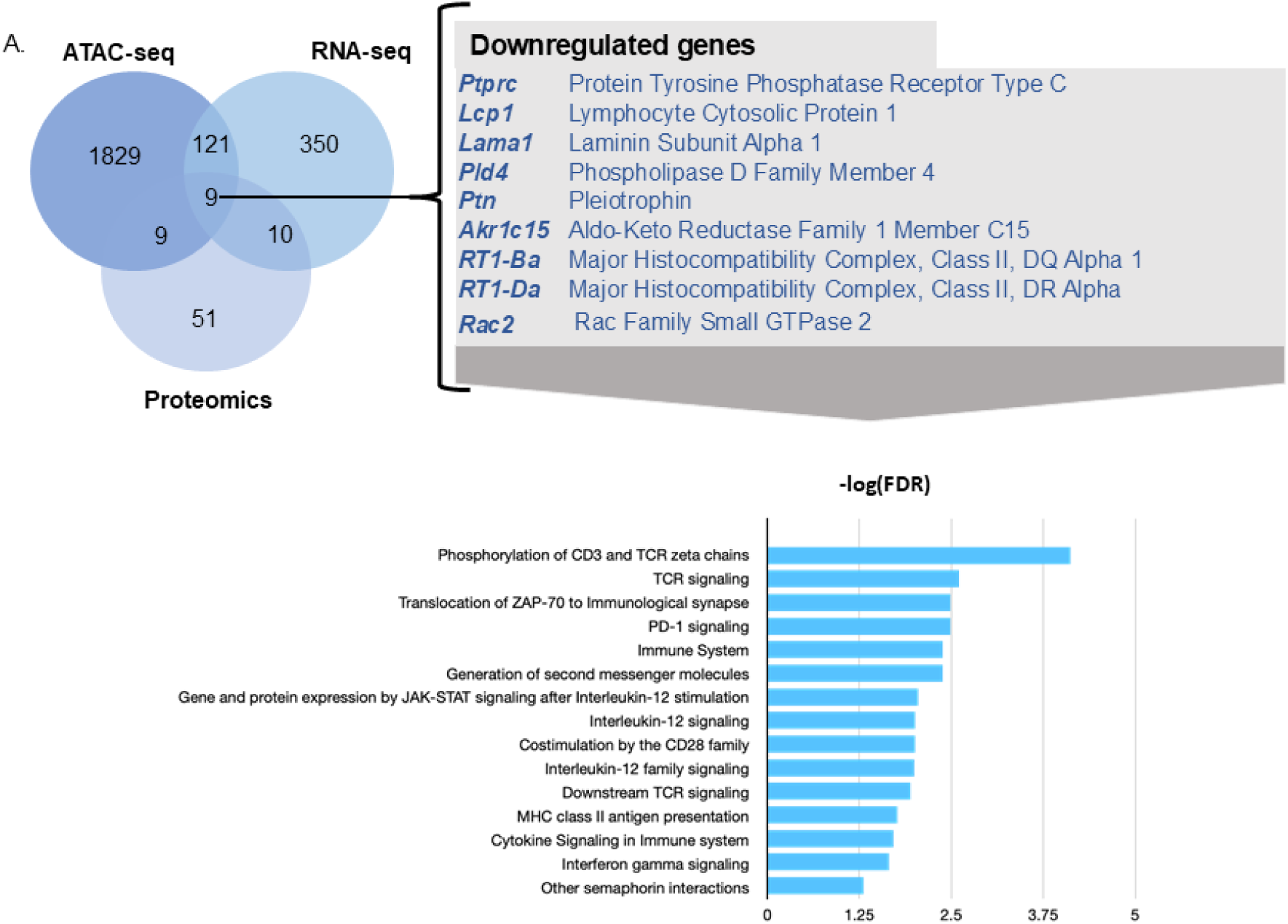
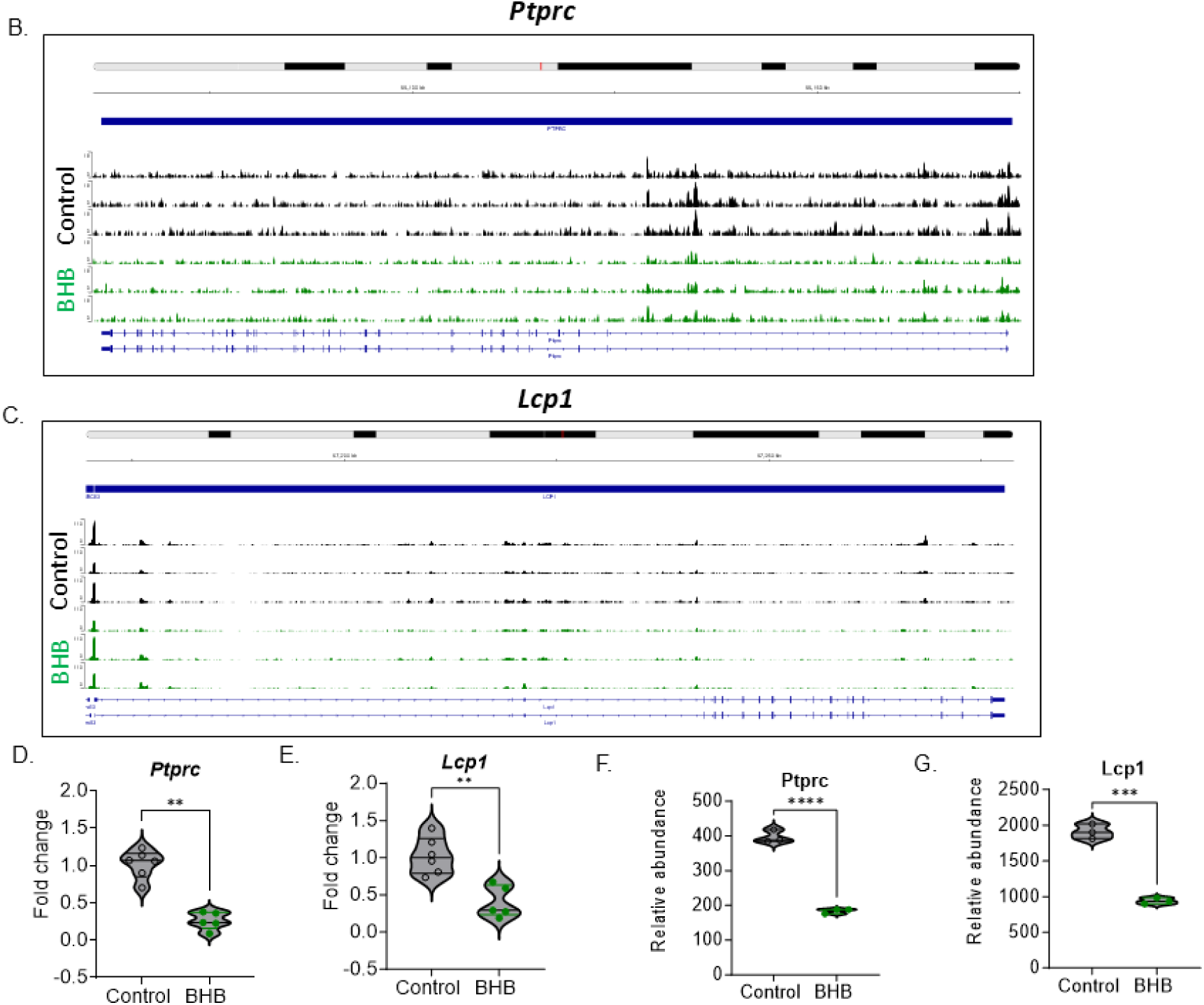
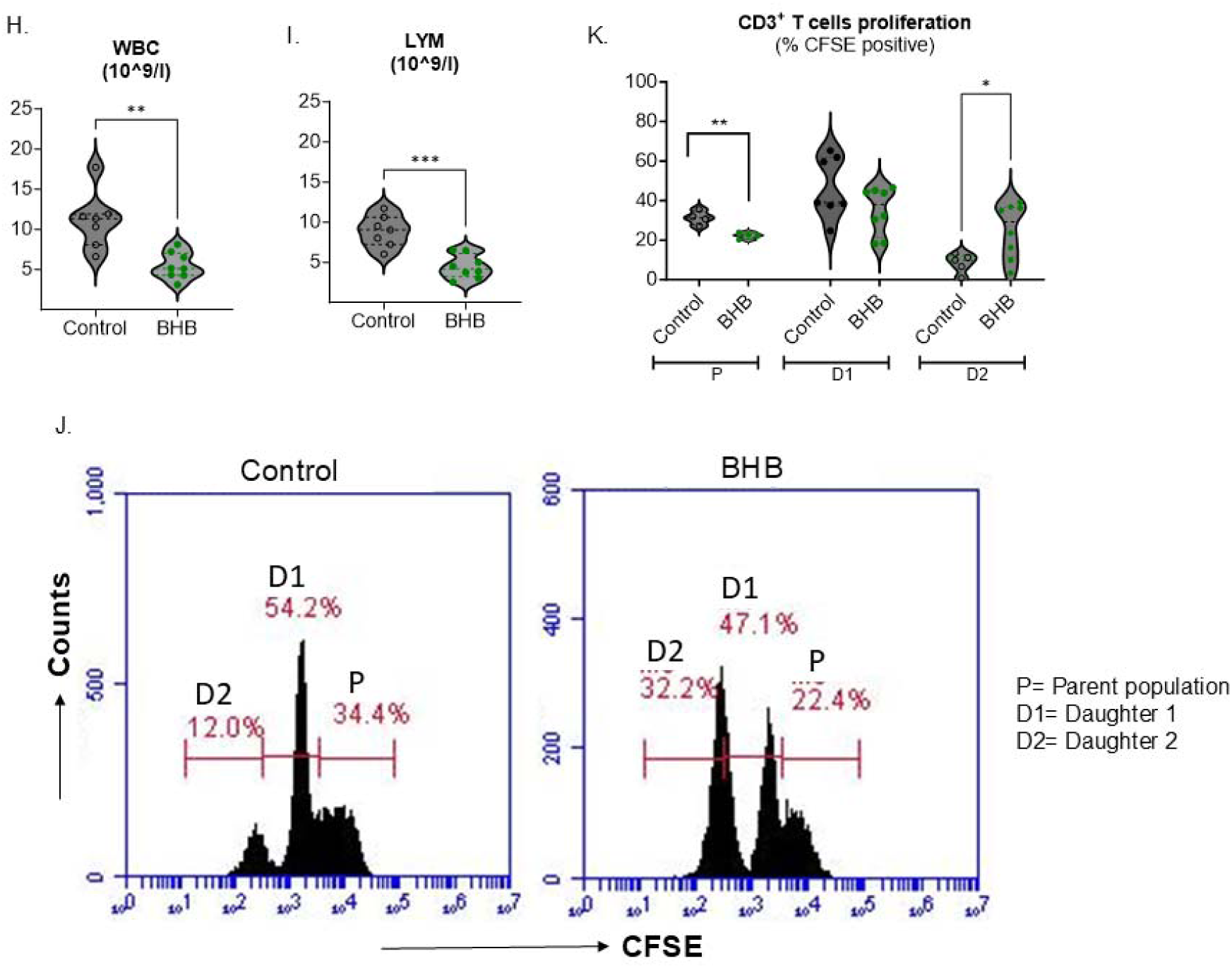
Chromatin accessibility, gene expression, protein levels, and CD3^+^ T cells proliferation were altered with 1,3-butanediol treatment. (A) Venn diagrams showing common downregulated genes and proteins leading to downregulation of immune function pathways. Different shades of blue circles are used to indicate ATACseq, RNAseq, and Proteomics data. The gray box lists the names of common downregulated genes and proteins, beneath which are shown the associated downregulated pathways. (B) Chromatin accessibility at the promoter region of *Ptprc* and (C) Chromatin accessibility at the *Lcp1* promoter. The y-axis represents chromatin cut sites and thus open chromatin, and x-axis represents the chromatin location of genes of interest. Black peaks represent the control group, and green peaks represent BHB group. (D-E) Real time PCR data showing reduced expression with *Ptprc* and *Lcp1* in the 1,3-butanediol supplementation. (F-G) Normalized relative abundances of Ptprc (Cd45) and Lcp1 detected in the quantitative proteomics study. Black open circle– control, green closed circle-BHB. (H-I) Complete blood count (H) Peripheral white blood cells and (I) Lymphocytes in the control and BHB treated male S rats. (J) Representative histogram for percent CFSE-positive CD3+ T cells. (K) Quantification for percent CFSE-positive CD3^+^ T cells in control and BHB groups after 5 days. Data represented as Mean ± SEM and N was plotted for control and BHB group. WBC: white blood cells, Lym: Lymphocytes, CFSE: carboxyfluorescein diacetate succinimidyl ester, CD3: Cluster of differentiation 3, P= Parent population, D1= Daughter 1, and D2= Daughter 2 population All data are mean±SEM, *p<0.05, **p< 0.01, ***p< 0.001 and ****p<0.0001.

### Upregulation of Acaa1b-mediated fatty acid oxidation increased peroxisomal biogenesis

Fatty acid oxidation occurs both in mitochondria and peroxisomes^22–24^. Whereas mitochondria are the main site of oxidation of medium-and long-chain fatty acids, peroxisomes catalyze the β-oxidation of a distinct set of fatty acids, including very-long-chain fatty acids^25–27^. While Hmgcs2 is located within mitochondria, interestingly, another target of histone β-hydroxybutyrylation, Acaa1b, which is responsible for β-oxidation of fatty acids, is located within peroxisomes^28,29^. In response to metabolic stress, peroxisomes proliferate by upregulating their biogenesis^30^. One of the factors required for peroxisomal biogenesis is peroxisomal biogenesis factor 11 γ^31^. Interestingly, proteomic data revealed that Pex11γ was increased in the BHB group, which indicated that peroxisomal biogenesis was promoted in the BHB group (Supplementary Figure 2A). To further confirm this observation, we examined the renal sections for the abundance of peroxisomes using the classical marker, peroxisomal membrane protein 70 (Pmp70). Peroxisome staining was more prominent in the BHB group compared to control (Supplementary Figure 2B and 3C) suggesting that the upregulation of the peroxisomal target of histone-β-hydroxybutyrylation, Acaa1b, placed an increased demand for fatty acid oxidation, which in turn promoted peroxisome biogenesis.

### β-hydroxybutyrylation-mediated enhanced fatty acid oxidation inhibits Mammalian Target of Rapamycin Complex 1 (mTORC1)

Stimulation of peroxisome biogenesis by drugs such as rapamycin are known to inhibit mammalian target of rapamycin complex 1 (mTORC1) activity^32,33^. Hyperactivation of mTORC1 signaling is associated with several human diseases whereas suppression of mTORC1 is known to curb senescence, extend lifespan of yeast, *C. elegans*, *Drosophila* and mice and promote autophagy^34^. Importantly, mTORC1 inhibits fatty acid β-oxidation, whereas inhibition of mTORC1 by rapamycin promotes fatty acid oxidation^35^. Importantly, rapamycin suppresses mTORC1 signaling to ameliorate kidney injury and hypertension in S rats^36^. Based on the similarity in function between rapamycin and the targets of β-hydroxybutyrylation to promote fatty acid oxidation, we hypothesized that mTORC1 is inhibited in the BHB group to protect kidneys. In support, mTORC1 was significantly inhibited in the BHB group compared to control (Figure 3D, 4E, Supplementary Figure 3A and 4B).

### Chromatin remodeling by histone β-hydroxybutyrylation resulted in downregulation of immune function genes

Next, we examined the downregulated loci, transcripts and proteins in the BHB group compared to control. Similar to the shared upregulated genes and proteins, the combinatorial analysis using ATAC-seq, transcriptomic, and proteomic datasets showed that 9 proteins were downregulated in all the 3 analyses (Figure 6A). Among these, the top 2 were protein tyrosine phosphatase receptor type C (*Ptprc*) and Lymphocyte cytosolic protein 1 (*Lcp1*). Chromatin accessibility of the promoter regions of *Ptprc* and *Lcp1* were lower in the BHB group compared to control (Figure 6B and Figure 6C). These results indicate that histone β-hydroxybutyrylation mediated chromatin compaction of the promoter regions of *Ptprc* and *Lcp1* contributed to the observed lower transcription (Figure 6D, 7E and Supplementary Figure 4A, 5B) and translation (Figure 6F and 7G) of these loci.

Unlike the upregulated pathways of energy metabolism, reactome pathway analysis of the downregulated genes showed that the overall downregulated pathways due to histone β-hydroxybutyrylation were related to immune function. These were phosphorylation of CD3 and TCR zeta chains, downstream TCR signaling, translocation of ZAP-70 to immunological synapse, PD-1 signaling, immune system, interferon gamma signaling, interleukin-12 family signaling, and TCR signaling (Figure 6A).

### Histone β-hydroxybutyrylation promoted downregulation of CD45 and LCP-1

Next, we focused on the two top genes which were prominently downregulated in our ATAC-seq study, *Ptprc* and *Lcp-1*. Lower expression of both *Ptprc* and *Lcp1* in the BHB group was confirmed by qPCR (Figure 6D and 7E). These data correlated with lower abundance of the protein products of these genes (CD45 and LCP-1, respectively) in the BHB group compared to the controls (Figure 6H and 7I). CD45 and LCP-1 are both important for the function of T cells and dysfunctional regulation of T cells are implicated in renal disease and hypertension. Therefore, we focused on examining T cells using CD3^+^ as a pan marker reporting for all T cells in our experimental groups.

### Lower expression of CD45 and LCP-1 enhanced CD3^+^ T cells proliferation

Both CD45 and LCP-1 are present in all hematopoietic cells^37–40^. Therefore, we examined if the downregulation of these two genes in the BHB group was facilitated by the epigenetic action of β-hydroxybutyrate reflected in immunomodulatory effects. Analysis of complete blood count (CBC) revealed a significant decrease in the level of circulating white blood cells (WBCs), specifically lymphocytes, in the BHB group (Figure 6H and 7I). Similarly, mean corpuscular volume, mean corpuscular hemoglobin, mean platelet volume, plateletcrit and platelet distribution width were also lower in the BHB group. These data affirm an immunomodulatory function of β-hydroxybutyrate (Figure 6A, Supplementary Figure 5A-6F). In alignment, the parent population of CD3+ T cells was lower in the BHB group compared to controls (Figure 6J). Intriguingly, CD3+ T cells from the BHB group exhibited increased proliferation as indicated by a higher population of dividing cells (D2) while maintaining a similar population of D1 populations (Figure 6J and 7K). These findings suggest that lower expression of CD45 and LCP-1 in the BHB group contributed to a rapid turnover of T lymphocytes by depleting existing parental cells and promoting the generation of a sufficient number of new effector T cells to maintain immune homeostasis.

## Discussion

This study was designed to delineate the molecular mechanism underlying the previously documented protective effect of BHB^9^. Specifically, we examined the known function of β-hydroxybutyrate to epigenetically remodel chromatin by histone β-hydroxybutyrylation^41^. Chromatin remodeling by histone β-hydroxybutyrylation exerted dynamic effects of distinctly modifying the renal transcriptome and proteome to promote both mitochondrial and peroxisomal regulation of fatty acid catabolism while parallelly dampening immune cell function. We have methodically dissected and characterized the major loci, transcripts and proteins contributing to the specific consequence of renal histone β-hydroxybutyrylation to conclude that epigenetic chromatin remodeling by histone β-hydroxybutyrylation simultaneously regulates the dual processes of upregulation of fatty acid catabolism and dampening of immune cells. These results constitute a new mechanism underlying the dynamic beneficial renoprotective effect of β-hydroxybutyrate.

Histone β-hydroxybutyrylation was first identified as a new epigenetic modification occurring in the mouse liver in response to starvation^41^, which is a condition simultaneously promoting ketosis and production of β-hydroxybutyrate^11,42,43^. Its epigenetic action to post translationally modify histones by β-hydroxybutyrylation has been subsequently studied in the context of a variety of pathologies including cardiomyopathy^44^, depression^45^, glomerulosclerosis^46^, and lung adenocarcinoma^47^, but none in any renal or hemodynamic studies. While our previous study on the benefit of β-hydroxybutyrate^9^ has been reproduced by others in different contexts such as preeclampsia and diabetic kidney disease, the underlying molecular mechanism has remained elusive^48–50^. In this context, our results presented in this study are the first to reveal that post translational modification of histones by β-hydroxybutyrylation is a novel mechanism protecting kidneys.

Pathway analysis revealed an upregulation of fatty acid metabolism. Mitochondria, a major organelle involved in fat metabolism, undergoes morphological changes based on nutrient availability and stress conditions. mTORC1 is a known regulator of numerous cellular processes, such as cell proliferation, metabolism, and cell growth^51^. mTORC1 regulates mitochondrial metabolism and controls mitochondrial biogenesis^52^. Mitochondrial fission and apoptosis are controlled by mechanistic/mammalian target of rapamycin complex 1 (mTORC1) mediated stimulated translation of mitochondrial fission process 1 (MTFP1)^53^. Our study showed a distinct mitochondrial morphological alteration occurring in response to β-hydroxybutyrate, whereby mitochondria were more circular. Reduced circularity of mitochondria is reported to increase fatty acid oxidation, which concur with our findings^54^. Further, our data demonstrate that increased fatty acid oxidation was linked to a significant inhibition of mTORC1. In support, rapamycin, which is a uniquely specific mTOR inhibitor, increases fatty acid oxidation^35^. Similarly, fenofibrate, which is reported to have beneficial cardiovascular and renal effects, increased fatty acid oxidation and inhibited mTOR^55^. Importantly, in the context of salt-sensitive hypertension and kidney injury, Kumar *et al*. (2017), have shown the inhibition of mammalian target of rapamycin 1 (mTORC1) ameliorates both hypertension and renal injury^52^. These data taken together with our findings reported in the current study leads us to propose that epigenetic chromatin remodeling by histone β-hydroxybutyrylation promotes the renal fatty acid oxidation-mTORC1 axis to protect kidneys and lower hypertension.

An interesting finding of our study is that Hmgcs2, which is the main enzyme catalyzing the production of β-hydroxybutyrate from Acetyl CoA, is also the target locus impacted by β-hydroxybutyrylation. This indicates that β-hydroxybutyrate autoregulates its own production by epigenetically remodeling chromatin to promote the transcription of Hmgcs2. If this process is perpetual, systemic β-hydroxybutyrate levels would remain consistently elevated after a single intervention to raise systemic β-hydroxybutyrate. Such a dysregulated continuous production of β-hydroxybutyrate would be detrimental as it would lead to ketoacidosis. Besides, β-hydroxybutyrate is elevated only during fasting and is lowered during the fed state^9^. Similarly, β-hydroxybutyrylation was discovered to be elevated only during fasting^41^. Collectively, these data support our conclusion that histone β-hydroxybutyrylation by β-hydroxybutyryrate is not perpetual but turned ‘off’ when β-hydroxybutyryrate is limiting and/or by yet undiscovered mechanisms.

A very low-carbohydrate diet is reported to enhance human T-cell immunity through immunometabolic reprogramming^56^. Such a low carbohydrate diet could promote ketosis and inhibit mTOR. mTOR inhibition by Rapamycin, which also lowers hypertension, is shown to promote T cell hypo responsiveness or anergy^52^. In alignment with these observations, the downregulated pathways generated from downregulated genes/proteins led to lowering of T cell signaling, which implies that the mechanism by which T cell signaling is affected by β-hydroxybutyrate is a result of epigenetic remodeling of chromatin to a condensed state in regions harboring *Ptprc* and *Lcp1*.

Overall, our findings reveal that renal histone β-hydroxybutyrylation causes a dynamic increase in chromatin accessibility to the promoter regions harboring lipid catabolizing genes while simultaneously decreasing accessibility to the promoter regions of genes contributing to immune-related functions. Together, these newly discovered mechanisms underly the beneficial effects of β-hydroxybutyrate on renal-hemodynamic health.

## Disclosure

All the authors declared no competing interests.

## Author Contributions

BJ designed the study, JM, SC, SA, IM, MX, BSY, BM, AK, RT, PS, WTG, VB, IDLS collected and analyzed data; JM, SC, MX, BSY, BM, AK, RT, PS, WTG, VB, IDLS, SA, IM, MVK interpreted the data, SA, IM, BJ, JM, SC, VB, WTG, IDLS, PS contributed to writing the manuscript, All authors read and edited the manuscript.

