## Supplemental File for "Ketone body mediated histone β-hydroxybutyrylation is reno-protective"

**Contact for Reagents and Resource Sharing.**

**Supplemental Methods**

**Experimental Models and Subject Details**

Female Dahl Salt-Sensitive (S) rats were less tolerant to the dose as indicated by >20% loss of body weight, whereby on the third week, 1, 3- Butanediol was lowered to (10% v/v). After 5 weeks of 1, 3- butanediol treatment, S rats were euthanized by thoracotomy and exsanguination via cardiac puncture under isoflurane anesthesia administered via a nose cone (5% in 100% O_2_). This was consistent with the American Veterinary Medical Association Guidelines for the Euthanasia of Animals. All euthanasia and tissue harvesting were performed in the University of Toledo Department of Laboratory Animal Resources. Serum and kidneys were collected and stored at -80ºC.

**Measurement of Serum BHB**

While the rats were under isoflurane anesthesia, but prior to thoracotomy and exsanguination by cardiac puncture, arterial blood was collected from the abdominal aorta in silicone-coated collection tubes specifically used for serum collection (BD Vacutainer). Blood was set aside to clot at room temperature for ~20 min, centrifuged at 2000 g for 15 min at 4°C and the serum collected was flash frozen in liquid nitrogen, and stored at -80°C.  Thawed serum was used to quantitate BHB using a colorimetric assay kit from Cayman Chemicals. Briefly, sera were diluted 1:5 using the assay buffer provided. Standards and samples were incubated with the developer solution at 25°C in the dark and optical density was measured at 450 nm using a SpectraMax M3 microplate reader from Molecular Devices (Sunnyvale, CA, USA).

**ATAC-Sequencing**

Library preparation: Flash frozen renal tissues were transported to the UM Epigenomics Core for ATAC-Seq library preparation. Briefly, the tissues were cryo-pulverized with a Covaris CryoPREP2 (Covaris,). Frozen tissue powder (100mg) was aliquoted and processed for isolation of nuclei, enriched using an Optiprep discontinuous gradient and tagmented according to the Omni-ATAC-Seq protocol^1^. The concentration of nuclei was assessed and a total of 50,000 nuclei were used for tagmentation. The final libraries were cleaned using Qiagen MinElute columns and AMPure XP beads before quantitation with the Qubit HS dsDNA kit, and quality assessment on a TapeStation HS D1000 kit. The libraries were pooled and quantitated by qPCR using KAPA’s Illumina Library quantitation kit before PE-150 sequencing on a NovaSeq6000 S4 at the University of Michigan advanced Genomics Core.

Pipeline: Data preprocessing and quality controlling were performed using the bioinformatics pipeline, snakemake^2^. FastQC (FastQC) (v0.11.8) was used for Quality control of raw reads and trimmed reads^3^. TrimGalore (TrimGalore) (v0.4.5) and cutadapt (Martin) (v1.15) were used for adapter trimming and filtering of low-quality bases. Trimmed reads were aligned to rn6 with Bowtie2^4^. Alignment quality control was performed using samtools^5^. Reads completely overlapping blacklisted regions (ENCODE Blacklist Regions) were removed with bedtools^6^. Sample-wise peaks were called with macs2^7^. Peaks over all samples were merged with bedops^8^ for principal component analysis and unsupervised clustering to assess the similarity of samples. Finally, a report was generated by MultiQC^9^. For ATAC specific QC metrics, ataqv (Ataqv) was used^10^.

Differential Peaks:

For each model and contrast, the edgeR R Bioconductor package^11^ (v3.28.1) was used to identify regions of differentially open chromatin (DOC). For each sample, the number of reads in the merged peaks is counted for each sample, and a library size normalization factor was determined. With no replicates, the BCV (biological coefficient of variance) parameter was tuned. Each model was then fitted using the glmFit function, and each contrast was tested with a likelihood ratio test and for the replicates, the common, trended, and tagwise negative binomial dispersions were calculated. Each model was then fitted using the glmQLFit function, and each contrast was tested with an empirical Bayes quasi-likelihood F-test. The DOC were then annotated to genic and CpG island annotations using the annotatr Bioconductor package (v1.12.1)^12^. The threshold used to determine DOC was false discovery rate (FDR) < 0.05 and fold change (FC) >1.

**RNA Extraction and Real Time PCR Analysis**

Total RNA was extracted from control and 1, 3-Butanediol fed rat kidneys using Trizol (Invitrogen). RNA was reverse transcribed to cDNA using SuperScript III kit (Invitrogen). Levels of mRNA expression of *Acaa1b*, *Hmgcs2*, *Cyp2d4*, *Cyp2e1*, *Ptprc* and *Lcp1* (primer sequence shown in Table S2) were analyzed by real-time PCR (Applied Biosystems), and expression levels relative to the expression of *B-Actin* and *L36a* were calculated by the 2^-ΔΔCt^ method^13^.

**RNA Sequencing**

RNA extracted with the Trizol method was assessed for RNA quantity and purity with a Bioanalyzer 2100 and RNA 6000 Nano LabChip Kit (Agilent, CA, USA, 5067-1511) and dispatched to LC Sciences (Houston, TX). RNA with RIN number > 7.0 were considered of high quality and used to construct sequencing library. Two rounds of mRNA purification were conducted from total RNA (5ug) using Dynabeads Oligo (dT) (Thermo Fisher, CA, USA). Following purification, the mRNA was fragmented into short fragments using divalent cations under elevated temperature (Magnesium RNA Fragmentation Module (NEB, cat.e6150, USA) under 94℃ 5-7min). Next, the cleaved RNA fragments were reverse transcribed to create the cDNA by SuperScript™ II Reverse Transcriptase (Invitrogen, cat. 1896649, USA), which were then used to synthesize U-labeled second-stranded DNAs with E. coli DNA polymerase I (NEB, cat.m0209, USA), RNase H (NEB, cat.m0297, USA) and dUTP Solution (Thermo Fisher, cat.R0133, USA). An A-base was then added to the blunt ends of each strand, preparing them for ligation to the indexed adapters. Each adapter contained a T-base overhang for ligating the adapter to the A-tailed fragmented DNA. Dual-index adapters were ligated to the fragments, and size selection was performed with AMPureXP beads. After the heat-labile UDG enzyme (NEB, cat.m0280, USA) treatment of the U-labeled second-stranded DNAs, the ligated products were amplified with PCR by the following conditions: initial denaturation at 95℃ for 3 min; 8 cycles of denaturation at 98℃ for 15 sec, annealing at 60℃ for 15 sec, and extension at 72℃ for 30 sec; and then final extension at 72℃ for 5 min. The average insert size for the final cDNA librarys were 300±50 bp. At last, we performed the 2×150bp paired-end sequencing (PE150) on an Illumina Novaseq™ 6000 following the vendor's recommended protocol. The sequence quality was verified using FastQC (<http://www.bioinformatics.babraham.ac.uk/projects/fastqc/>, 0.11.9) ^14,15^.

**Quantitative Proteomic Analysis by Mass-spectrometry**

Kidneys were lysed using RIPA Lysis and Extraction buffer according to the manufacturer’s instructions. Proteins were quantified using the BCA assay, diluted to. 2mg/ml and shipped to the University of Michigan Proteomics Resource Facility for proteomic analysis. The protocol for Tandem Mass Tag labeling and offline fractionation was followed as described in Cheng et al., 2020^16^. Briefly, the TMT 6plex kit (#90061) was used for TMT labeling according to the manufacturer’s protocol (ThermoFisher Scientific). 100 ug protein was taken from each respective groups and DTT was used for the reduction reaction and alkylation with 2 chloroacetamide was done after that. Precipitation of proteins was done using ice-cold acetone and overnight incubation at -20 °C. Precipitated protein pellet was resuspended in Triethylammonium bicarbonate (TEAB) and overnight digestion was carried out using trypsin. (Promega, V5113). Digested peptides were transferred to TMT reagents reconstituted by anhydrous acetonitrile. Hydroxylamine was added and incubated. All samples were combined and dried. Prior to MS analysis, high pH reverse phase fractionation was performed, and fractions were dried and reconstituted in loading buffer containing 0.1% formic acid and 2% acetonitrile. Liquid Chromatography-mass spectrometry analysis (LC-Multinotch MS3) was performed followed by using Proteome Discoverer (v2.1; Thermo Fisher) for data analysis. Identified proteins and peptides were filtered to retain only those that passed ≤1% FDR threshold. Quantitation was performed using high-quality MS3 spectra (Average signal-to-noise ratio of 10, less than 30% isolation inference, and data was normalized against total peptide).

**ChIP Assay**

Rat tissue was ground to a fine powder with a mortar and pestle then crosslinked with 1% formaldehyde and homogenized as in^17,18^. Nuclei were isolated^19^ in Lysis Buffer 1 (50mM HEPES-KOH, pH 7.5, 140mM NaCl, 1mM EDTA, 10% glycerol, 0.5% NP40, 0.25% TritonX, and protease inhibitors) then washed with Lysis Buffer 2 (10mM Tris-HCl, pH 8.0, 200mM NaCl, 1mM EDTA, 0.5mM EGTA and protease inhibitors). Nuclei were re-suspended in Lysis Buffer 3 (10mM Tris-HCl, pH 8.0. 100mM NaCl, 9mM EDTA, 0.5mM EGTA, 0.1% Na-Deoxycholate, and 0.5% N-lauryolsarcosine, protease inhibitors) and sonicated in 30 second pulses at 60% power, for a total of 6 minutes to get an average fragment size of 200-1000 base pairs. A small portion (1%) of the chromatin was saved as “Input”. Approximately 25µg of chromatin was pre-cleared for one hour at 4^o^C with Protein A Sepharose (Cytiva, Marlborough, MA, USA), then incubated overnight with an antibody to β-hydroxybutyryl-histone H3 (Lys 9) (H3K9BHB) (PTM Biolabs, Chicago, Il, USA) or normal rabbit IgG (Millipore Sigma, St. Louis, MO, USA) in Lysis Buffer 3, supplemented with 1.1% Triton X-100. Protein A sepharose was added the next morning and incubation was allowed to proceed for an additional four hours. The precipitated chromatin was then washed five times in RIPA buffer (50 mM HEPES, 500 mM LiCl, 0.1 mM EDTA, 1.0% NP-40 and 0.7% Na-Deoxycholate) and once with TE containing 50mM NaCl. Complexes were then eluted with 50 mM Tris [pH 8.0], 10 mM EDTA and 1.0% SDS. Crosslinks were reversed by overnight incubation at 65^0^C. After proteinase K digestion, DNA was purified by phenol/chloroform extraction and ethanol precipitation. Genomic regions of interest were identified from ATAC-seq data, and the sequences retrieved using the UCSC Genome Browser. Primers were designed using the Primer 3.0 tool version 4.1.0^20,21^. PCR amplifications were performed on an ABI 7500 system (Thermo Fisher, Waltham, MA, USA) using SYBR green master mix (Qiagen, Hilden, Germany). Data were analyzed using ABI 7500 software to generate cycle thresholds (CT values). Delta CT values were calculated by subtracting the CT values of the ChIPs obtained with the antibodies of interest from the CT values of the Input controls. The 2^-ΔΔCt^ method was then used to calculate enrichment of immunoprecipitated material. Average ChIP enrichment was calculated from three biological replicates, each amplified as three technical replicates. Primer sequences for ChIPs PCR are mentioned in supplemental table 2.

pathways^22^.

**Transmission Electron Microscopy and Analysis of Mitochondrial Morphology**

Fresh minced or formalin fixed tissues (not frozen) were placed into sodium cacodylate buffered with 3% glutaraldehyde (pH 7.4) at room temperature for 1 1/2 - 2 hours. Specimens were gently agitated throughout the tissue processing procedure. Samples were subsequently washed with 2-3 changes of fresh cacodylate buffer (pH 7.4) for a total time of 20-30 minutes, then immersed in cold (4ºC) 1% osmium tetroxide (OsO4) in cacodylate buffer (pH 7.4) for 1 1/2 - 2 hours at room temperature. Following OsO4 fixation 2-3 changes of fresh cacodylate buffer were used for washing for a total of 20-30 minutes. Tertiary fixation was included using an aqueous saturated solution of uranyl acetate in distilled water (pH 3.3) for 1.0 hour. The tissues were dehydrated at room temperature with an increasing concentration of ethanol (50%, 70%, 90%, 95%, and 100%) followed by 100% acetone. Samples were then infiltrated with a 50% acetone**:** 50% embedding media (Araldite 502/Embed 812) overnight at room temperature. The next day, tissues were embedded in appropriately labeled embedding capsules filled with fresh 100% Araldite 502/Embed 812 embedding media. Capsules were polymerized in a 65ºC oven overnight. A Leica Reichert Ultracut S Ultramicrotome (Austria) was used to obtain 75-80 nm thin sections that were collected upon 300 mesh copper grids and post-section stained with uranyl acetate and Reynolds lead citrate and examined using a Talos L120C G2 3D Tomography Transmission Electron Microscope (ThermoFisher Scientific, USA).  Images were obtained at 6700X magnification to provide a minimum of 1000 mitochondria per group for image analysis to calculate average mitochondrial area, perimeter and mitochondrial circularity index using ImageJ software from NIH.

**Supplemental Table 1: Key Resources Table**

| **REAGENT or RESOURCE** | **SOURCE** | **IDENTIFIER** |
| --- | --- | --- |
| **Antibodies** | | |
| Anti-β-hydroxybutyryl-Histone H3 (Lys9) rabbit pAb | PTM Bio | Cat# PTM-1250 |
| Anti-Histone H3 rabbit pAB (CT) | PTM Bio | Cat# PTM-1002 |
| Rabbit IgG | Millipore Sigma |  |
| Phospho-S6 Ribosomal Protein (Ser235/236) Rabbit mAb | Cell Signaling Technology | Cat# 4858 |
| S6 Ribosomal Protein Rabbit mAb | Cell Signaling Technology | Cat# 2217 |
| **Chemicals, Peptides and Recombinant Proteins** | | |
| (±)-1,3-Butanediol | Sigma-Aldrich | Cat# B84785 |
| Trizol reagent | Invitrogen | Cat# 15596-026 |
| RIPA Lysis buffer | Thermofisher Scientific | Cat#89900 |
| **Critical Commercial Assays** | | |
| β-Hydroxybutyrate Colorimetric Assay kit | Cayman Chemicals | Cat# 700190 |
| **Deposited Data** | | |
| **Experimental Models: Organisms/**  **Strains** |  |  |
| Dahl Salt sensitive rats/ S | University of Toledo | https://rgd.mcw.edu/rgdweb/report/strain/  main.html?id=69369 |
| SS-*Nlrp3^em2^*rat | Medical College of Wisconsin | MCW Gene Editing Rat Resource Center |
| **Oligonucleotides** |  |  |
| See Supplementary Tables 1 and 2 for primer sequences | Current manuscript | N/A |
| **Software and Algorithms** | | |
| GraphPad Prism 10.1.0 | GraphPad software | https://www.graphpad.com/ |
| ShinyGO 0.77 | South Dakota State University | http://bioinformatics.sdstate.edu/go/ |
| Reactome v86 | Reactome | https://reactome.org/ |
| SnapGene 7.0 | Dotmatics | https://www.snapgene.com/ |
| ImageJ | National Institute of Health | https://imagej.net/ij/ |
| Quantstudio Design & Analysis | Thermo Scientific | https://apps.thermofisher.com/apps/da2/#/home/ |
| Microsoft 365 | Microsoft | https://www.office.com/ |
| **Reagents and Resources** | | |
| Harlan Teklad low salt diet | Envigo | Cat#TD7034 |
| Harlan Teklad high salt diet | Envigo | Cat#TD94217 |
| Radiotelemetry transmitters | Data Science Int. | PA-C10 |
| SpectraMax M3 | Molecular Devices | N/A |
| Nanodrop 2000 | Thermo Scientific | N/A |
| VS120 Virtual Slide Microscope | Olympus | N/A |
| Talos L120C G2 3D Tomography Transmission Electron Microscope | ThermoFisher Scientific | N/A |
| Covaris CryoPREP2 | Covaris | N/A |
| NovaSeq6000 Sequencing system | Illumina | N/A |

**Supplemental Table 2: Primers for Real Time q-PCR**

| **Gene** | **Sequence (5’-3’)** (Forward & Reverse Primers) |
| --- | --- |
| *Hmgcs2* | CGGAGACGCATGTCCCCTGA |
|  | AATGGTTGTATGGATTGGCCTCCTT |
| *Ptprc* | ACCTGCTCGCACCACTGAAT |
|  | GCAGAGGGGGTCTGTGAGTC |
| *Cyp2e1* | CCTGCATGGCTACAAGGCTG |
|  | GGGAAAACCTCCGCACATCC |
| *Cyp2d4* | ATGATGAGAACCTGCGTGTGG |
|  | TCGATTTCCTGTTGTACTCGGC |
| *Acaa1b* | CTTCTGTCGGCCGTGTTGAC |
|  | GGCCCCGGGCTGAAG |
| *Lcp1* | AGGGATCTGTGCGATCGGTG |
|  | TCGGGTTCATGGGGATGACG |
| *L36a* | ATTGTGCTCAGGCTTGAGTGT |
|  | TGG ATCACTTGACCCTTT |

**Supplemental Table 3: Primers for ChIP PCR**

| **Gene** | **Sequence (5’-3’)** (Forward & Reverse Primers) |
| --- | --- |
| *Hmgcs2* | ACCTTTGGCCCAGTTTTTCT |
|  | GGTGAGAGCGAAGATTCCTG |
| *Ptprc* | AAGCCCTCCTGTGTCCAATA |
|  | GTGCACATGTGTAGGGGTGT |
| *Cyp2e1* | TAAAGGCTCCAAGGTTCAGC |
|  | GAGAAGGAGGGTGGCTCATC |
| *Cyp2d4* | GGGAGAGGTGAAGGAGGAGA |
|  | GGATCTGCCTCCTCAGAGTG |
| *Acaa1b* | CCTAGCATCGATGTGTGCAG |
|  | AGAGCCCAAAACTCGAAAGG |
| *Lcp1* | CAGAAAGGCCAAGGAGTCAG |
|  | CCACCAATAACGGAGATGCT |

**Supplementary Figure Legends**

**Supplementary Figure 1: BHB supplementation downregulates pathway related to immune cell function.** (A) Pathways regulated by downregulated genes within the BHB group, (B) Pathways regulated by downregulated proteins in BHB group.

**Supplementary Figure 2: Validation of common upregulated and downregulated genes in female rat kidney.** (A-D) Real time PCR data depicting significant upregulation of *Hmgcs2, Acaa1b*, *Cyp2d4* in the BHB supplemented female rats. (E-F) *Ptprc* and *Lcp1* showed downregulated trend in the BHB treated group compared to the control. Housekeeping gene: β-actin. All data are mean±SEM, *p< 0.05 and ***p< 0.001. ns: Data trending but not significant.

**Supplementary Figure 3:** **Lower protein cast and higher peroxisome in rats treated with BHB.** (A) Proteomics data depicting upregulation of Pex11γ level in BHB supplemented male rat kidney. Data are mean± SEM, *p< 0.05. (B-C) Representative images (8x and 40x magnifications) of kidney sections with immunostaining for peroxisomes (Pmp70, brown color); (B) control S rats and (C) 1,3-butanediol fed rats. Relatively higher intensity of brown color was observed in the BHB group compared to control. IHC-PMP: Immunohistochemistry-peroxisomal membrane protein.

**Supplementary Figure 4: BHB inhibits MTORC1 activity in the BHB supplemented female kidney.** (A-B) Reduced phospho-S6/S6 ribosomal protein in female kidneys from the BHB group compared to controls. Data are mean ±SEM, *p< 0.05.

**Supplementary Figure 5: Chromatin Immunoprecipitation assay, food intake and activity in male rats.** (A) Comparable Chromatin immunoprecipitation data from kidneys of male rats for *Ptprc* and *Lcp1* promoters with H3K9 β-hydroxybutyrylated histones. (n=3 replicates/group) (B) Significantly lower 24-hour food intake in the BHB group. (n=5/group) (C) No differences in 24-hour average activity between BHB and control groups. All data are mean ± SEM; Ns: p>0.05, ***p< 0.001.

**Supplemental Figure 6**: BHB restores blood parameters as indicated by the complete blood count. (A) Mean corpuscular volume (MCV). (B) Mean corpuscular hemoglobin (MCH). (C) Mean platelet volume (MPV). (D) Plateletcrit (PCT). (E-F) Platelet distribution width (PDW). Data represented as Mean ± SEM. *p<0.05, **p<0.01, ***p<0.0001.

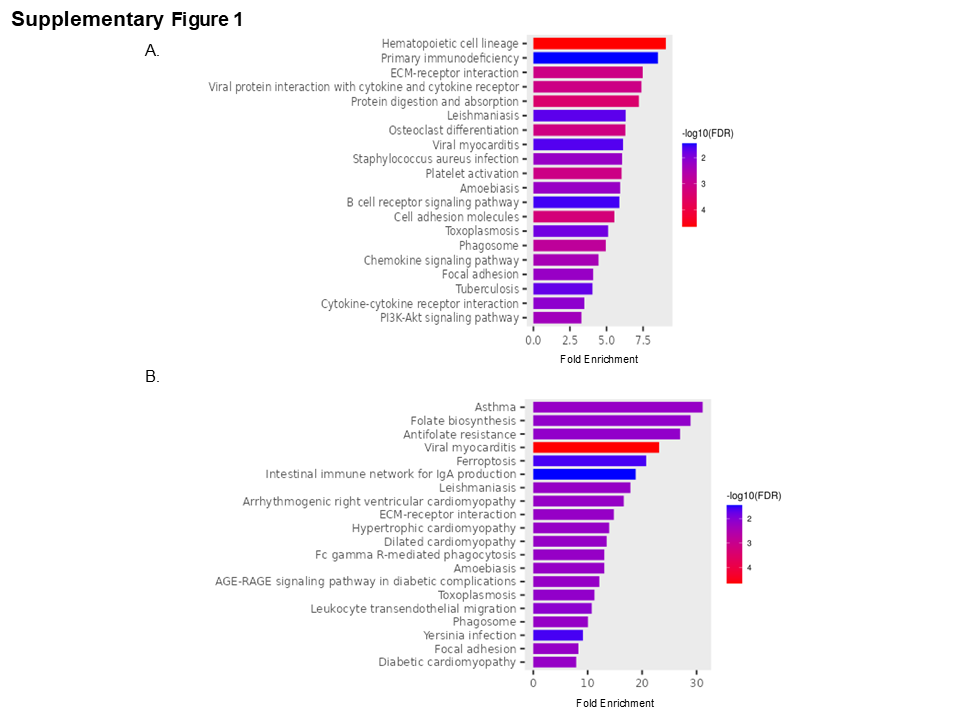

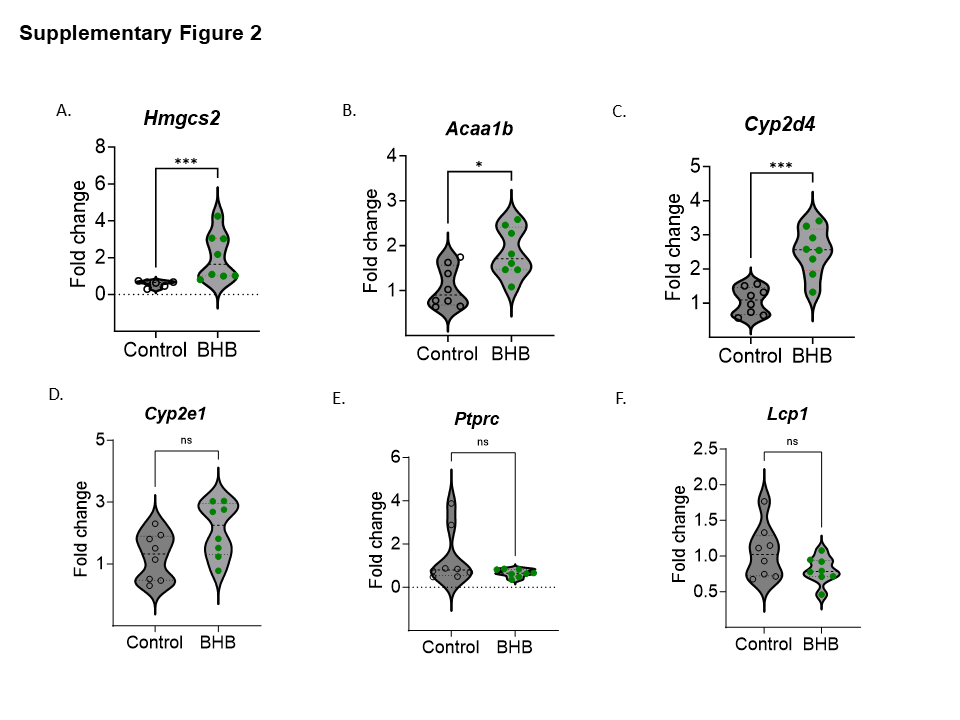

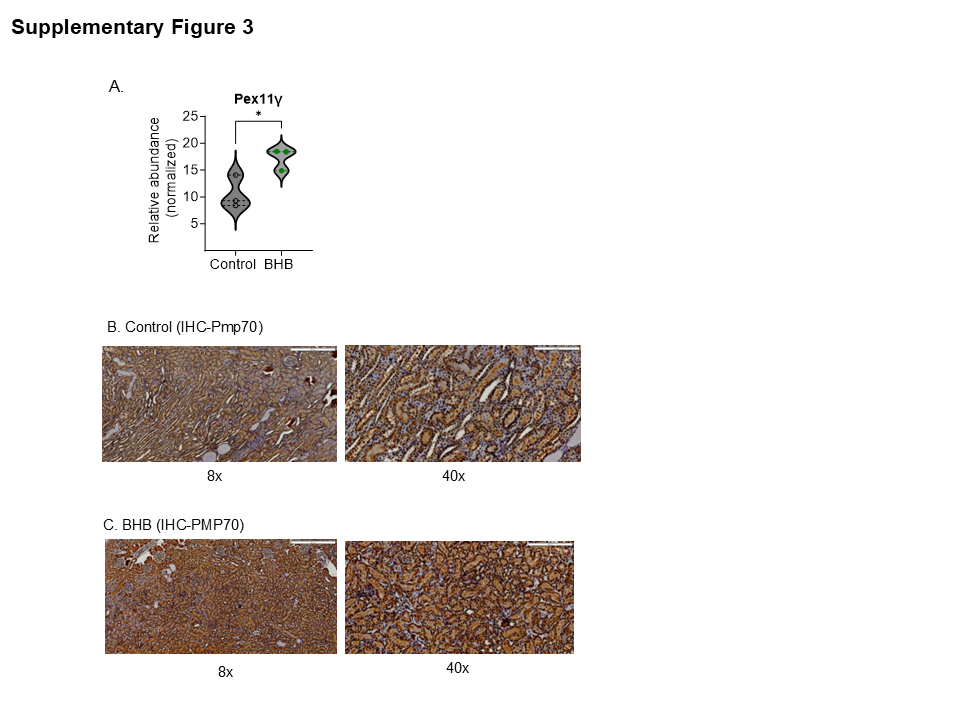

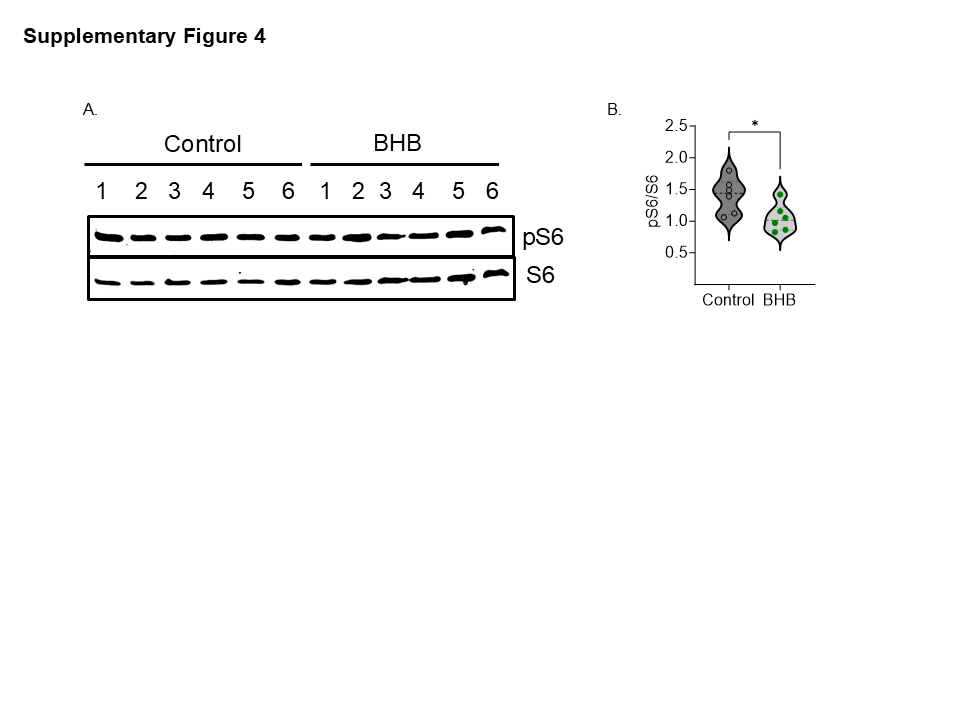

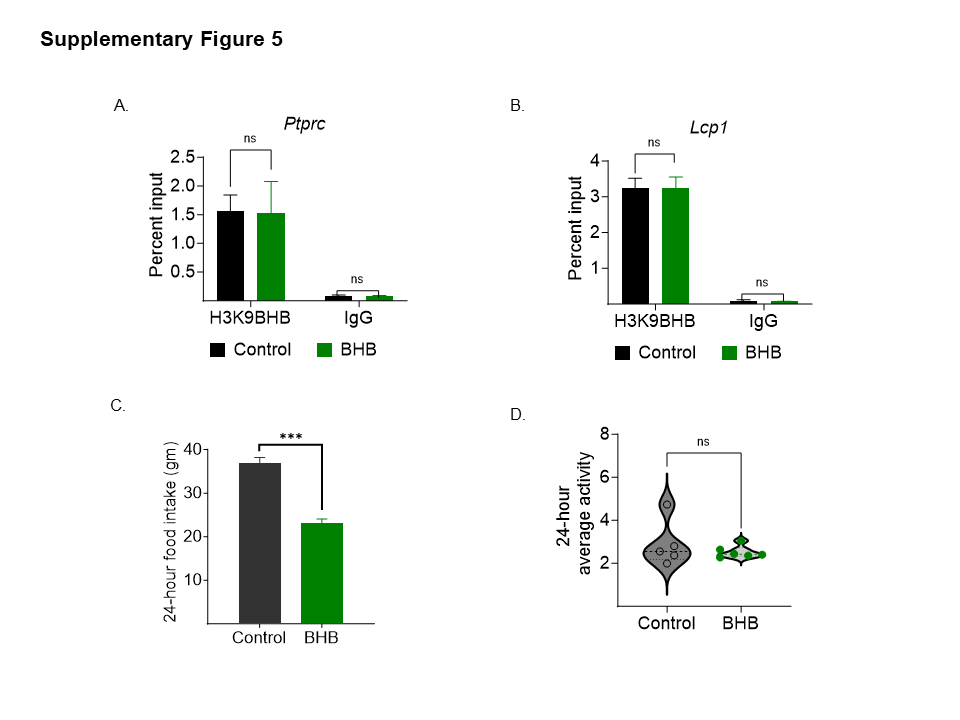

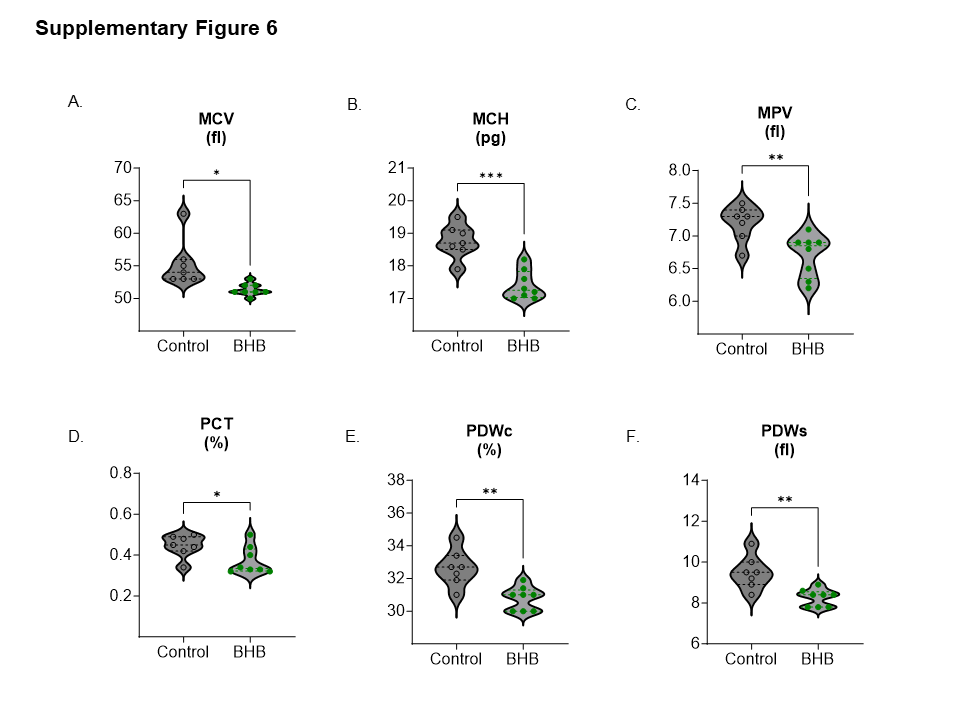
